# Chronically elevated corticosterone impairs dopaminergic transmission in the dorsomedial striatum by sex-divergent mechanisms

**DOI:** 10.1101/2022.10.03.510627

**Authors:** Ashley L. Holloway, Michael D. Schaid, Talia N. Lerner

## Abstract

**Background:** Major depressive disorder (MDD) is a leading cause of disability worldwide. Individuals with MDD exhibit decreased motivation and deficits in reward processing along with chronically elevated levels of the ‘stress hormone,’ cortisol. However, the mechanistic relationship between chronically elevated cortisol and behavioral deficits in motivation and reward processing remains unclear. Given that women are diagnosed with MDD at twice the rate of men, it is important to understand whether the mechanisms linking chronically elevated cortisol to the symptoms of MDD differ by sex.

**Methods:** We used subcutaneous implants to chronically elevate free plasma corticosterone (rodent homolog of cortisol; ‘CORT’) in male and female mice and examined changes in behavior and dopamine system function. We used operant training to assay reward-guided motivation and high-performance liquid chromatography to quantify striatal dopamine content. We assessed dopamine transporter (DAT) function using *ex vivo* slice imaging and *in vivo* fiber photometry in combination with a fluorescent dopamine sensor, dLight1.3b. We further quantified DAT expression and phosphorylation using western blot.

**Results:** We found that chronic CORT treatment impaired reward-seeking in both sexes. In female but not male mice, CORT treatment reduced dopamine content in the dorsomedial striatum (DMS). In male but not female mice, the function of the dopamine transporter (DAT) was impaired in DMS.

**Conclusions:** Chronic elevation of CORT impairs reward-seeking by impairing dopaminergic transmission in the DMS, but via different mechanisms in male and female mice. A better understanding of these sex-specific mechanisms could lead to new directions in MDD diagnosis and treatment.

## Introduction

Major depressive disorder (MDD) is a leading cause of disability worldwide, affecting an estimated 5% of adults (1). Individuals with MDD exhibit decreased motivation and deficits in reward processing (2,3). One important factor that precipitates and exacerbates MDD is stress (4). CORT (corticosterone in rodents, cortisol in humans) is the body’s primary stress hormone, released by the adrenal gland both in a regular circadian rhythm and in response to stressful events. In individuals with MDD, normal rhythms of high and low CORT are flattened, leading to dysregulated and chronically elevated levels of CORT (5). Despite decades of research, it remains unclear how chronically elevated CORT contributes to MDD symptomology.

In rodent preclinical models, chronic elevation of circulating CORT impairs operant responding for rewards, suggesting that elevated CORT may also be responsible for impaired reward processing in humans (6,7). However, rodent studies have only been carried out in males, leaving open the question of sex differences in the effects of chronically elevated CORT. Given that MDD is twice as common in women vs men, sex differences in biological responses to chronically elevated CORT are important to assess. Furthermore, the biological mechanisms underlying CORT-induced impairments in operant responding in either sex remain unclear.

We hypothesized that chronically elevated CORT could impact operant responding by altering dopaminergic transmission. Dopaminergic transmission in the striatum regulates reward processing (8). Specifically, dopamine in the nucleus accumbens core (NAcc) is critical for effortful responding for rewards, while dopamine in the dorsomedial striatum (DMS) is crucial for action-outcome learning and behavioral flexibility (9–12). Dopaminergic transmission within the striatum occurs in two modes: tonic and phasic (13,14). Tonic dopamine is the sustained level of extracellular dopamine in the striatum. It arises from the regular, tonic firing activity of dopamine neurons and is also tightly regulated by the kinetics of dopamine reuptake into terminals by the dopamine transporter, DAT (15,16). Tonic dopamine is hypothesized to govern motivation (17,18). Phasic dopamine transmission occurs in discrete epochs on top of tonic dopamine and is controlled by bursting activity of dopamine neurons. Phasic dopamine transmission invigorates responding for rewards and facilitates associative learning about cues and actions that lead to rewards (19–22). We examined whether impaired operant responding for rewards following chronic CORT treatment was associated with impaired tonic and phasic dopaminergic transmission in the NAcc and DMS of male and female mice.

## Methods

### Animals & Housing

Adult (10+ weeks) male and female C57BL6/J mice were used for all experiments. Mice were group housed by sex and treatment, and had ad libitum access to food and water, unless otherwise specified. Mice were housed on a 14:10 hour light/dark cycle, in a temperature- and humidity-controlled environment. All experimental procedures were approved by the Northwestern University Animal Care and Use Committee. All experiments were completed at zeitgeber time 4-6 (4-6 hours after lights-on).

### Subcutaneous Pellet Implants

At 10+ weeks of age, mice were anesthetized with isoflurane and given analgesics to minimize pain after surgery. Hair was removed from the lateral portion of the neck using Nair, and the skin was swabbed with alcohol and iodine. A small incision was made, and Placebo or Corticosterone (35 mg; 60-day release; Innovative Research of America) slow-release pellets were implanted subcutaneously in the space between the shoulder and neck. The incision was closed with non-absorbable sutures. For *ex vivo* slice imaging and *in vivo* photometry experiments, pellets were implanted during stereotaxic surgeries.

### Stereotaxic Surgeries

At 10+ weeks of age, mice were anesthetized with isoflurane and given analgesics to minimize pain. Hair on the skin of the top of the head was removed using Nair, then swabbed with alcohol and iodine. A single incision was made down the midline of the skull, then a hole was drilled above the injection site for the dorsomedial striatum (DMS; +0.8 A/P, 1.5 M/L, -2.8 D/V, relative to bregma) and nucleus accumbens core (NAcc: +1.6 A/P, 0.8 M/L, -4.1 D/V). 500 nL of AAV9-CAG-dLight1.3b (7×10_11_ VG/mL) (23) was injected into the DMS and NAcc at a rate of 100 nL/min using a Hamilton syringe. The needle was left in place for five minutes after injection before being slowly retracted. For fiber photometry experiments, a fiber optic (Doric, 400 µm core, 0.66 NA) was implanted over the DMS injection site. The hemispheres of injection sites were counterbalanced across treatment groups and sexes.

### Operant Conditioning

Mice were food restricted to 85% their ad libitum weight and were monitored for maintenance of this weight throughout operant training. All operant sessions lasted 60 minutes, or until mice received the maximum number of rewards available (50 rewards). Mice were initially trained to acquire sucrose rewards from the reward port of an operant box (Med Associates) in the absence of any contingency. Mice were then advanced to a fixed-ratio (FR) schedule of training during which they had to nosepoke once for one sucrose pellet (FR-1). After earning at least 30 rewards for two consecutive days (criterion for advancement), mice were advanced to FR-3 training, in which they had to nosepoke three times for one sucrose pellet. After reaching criterion for advancement, mice were advanced to FR-5 training.

### High Performance Liquid Chromatography and Electrochemical Detection of Dopamine

Biogenic amines were measured in the Vanderbilt University Neurochemistry Core.

### Ex vivo dLight1.3b Imaging

At least 4 weeks after pellet implantation and stereotaxic surgery to inject AAV9-CAG-dLight1.3b in the DMS and NAcc, mice were anesthetized with Euthasol (Virbac, 1 mg/kg) and transcardially perfused with ice-cold N-methyl-D-glucamine (NMDG) (24) artificial cerebrospinal fluid (ACSF) containing (in mM): 92 NMDG, 2.5 KCl, 1.2 NaH2PO4, 30 NaHCO3, 20 HEPES, 25 Glucose, 5 Na-Ascorbate, 2 Thiourea, 3 Na-Pyruvate, 10 MgSO4, 0.5 CaCl2. Coronal tissue sections (300 µm thick) containing the DMS were cut using a vibratome (Leica VT1200) and transferred to NMDG ACSF at 33°C. Slices were incubated for 15 minutes before being transferred to 33°C HEPES ACSF containing (in mM): 92 NaCl, 2.5 KCl, 1.2 NaH2PO4, 30 NaHCO3, 20 HEPES, 25 Glucose, 5 Na-Ascorbate, 2 Thiourea, 3 Na-Pyruvate, 1 MgSO4, 2 CaCl2. Slices were then transferred to HEPES ACSF at room temperature for 15 minutes before being held in recording ACSF containing (in mM): 125 NaCl, 26 NaHCO3, 1.25 NaH2PO4, 2.5 KCl, 1 MgCl2, 2 CaCl2, 11 Glucose. All solutions used were saturated with carbogen (95% Oxygen, 5% Carbon Dioxide) and their pH and osmolarity were adjusted to 7.3-7.4 and 300±5 mOsm, respectively. Slices were transferred to a recording chamber in ACSF, held at 30-32°C. For recording, ACSF contained blockers for AMPARs (NBQX, 5µM), NMDARs (D-AP5, 50µM), nAChRs (DHβE, 1µM), GABA_A_Rs (Picrotoxin, 50µM), and GABA_B_Rs (CGP-54626, 2µM). Dopamine release was evoked using a bipolar stimulating electrode (FHC, Inc.) placed ∼300 microns from the imaging site. All stimulations were 4V, with a pulse width of 0.5ms. After baseline recordings, the DAT inhibitor GBR-12909 (1µM) was applied to slices, followed by the OCT3 inhibitor Normetanephrine (50µM). dLight1.3b fluorescence was imaged using a scientific CMOS camera (Hamamatsu Orca-Flash 4.0LT), with a sampling rate of 33 Hz. dLight1.3b tau-off values were calculated using a custom MATLAB script.

### In vivo dLight1.3b Fiber Photometry

Fiber photometry experiments occurred at least four weeks after pellet implantation and stereotaxic surgeries. Mice were attached to a fiber optic patch cord (Doric, 400 µm core, 0.66 NA) and gently placed in an open field (28 × 28 cm). After 10 minutes in the open field, mice were injected with a DAT inhibitor, GBR-12909 (20 mg/kg, i.p.), and returned to the open field for another 40 minutes. All fiber photometry recordings were performed using a fiber photometry rig with optical components from Doric lenses controlled by a real-time processor from Tucker Davis Technologies (TDT; RZ5P). TDT Synapse software was used for data acquisition. 465nm and 405nm LEDs were modulated at 211 Hz and 330 Hz, respectively. LED currents were adjusted to return a voltage between 150-200mV for each signal, were offset by 5 mA, were demodulated using a 4 Hz lowpass frequency filter. Fiber photometry data was collected throughout the entire time that mice were in the open field. GuPPy, an open-source Python-based photometry data analysis pipeline, was used to determine dLight1.3b transient timepoints (25). A custom MATLAB script was used to calculate dLight1.3b area-under-the-curve (AUC). Briefly, dLight1.3b fluorescence was normalized by fitting it with the isosbestic fluorescence curve, filtered with a 1 second median filter, downsampled to 1Hz, then dLight1.3b AUC was calculated in one-minute bins for the entire trace, and normalized to the average 1-minute AUC of the drug-free period by subtraction (26). Locomotor activity was recorded using Noldus Ethovision XT 16.

### Western Blot

An equal amount of protein from each sample was loaded in a Tris-Glycine gel (Invitrogen). Protein was transferred to a PVDF membrane and blocked in either 5% bovine serum albumin (BSA) in Tris-buffered saline + 0.1% Tween-20 (TBS-T) for phospho-DAT, or 5% non-fat milk (NFM) in TBS-T for DAT and Beta-Actin. Membranes were blocked for one hour at room temperature, then incubated in primary antibody in blocking buffer overnight at 4°C. Membranes were washed in TBS-T, then incubated in secondary antibody in blocking buffer for 1-2 hours at room temperature. Membranes were imaged using a Licor Odyssey Fc Imaging System. Densitometric analysis was completed using ImageJ. Protein expression was normalized to the average of the sex-matched Placebo group for statistical analysis.

Additional methods are provided in the supplement.

## Results

### Corticosterone pellet implant chronically increases total plasma corticosterone in male mice and decreases plasma corticosteroid binding globulin levels in female mice

To chronically elevate corticosterone (CORT) levels in mice, we implanted male and female mice with a subcutaneous slow-release CORT pellet (35 mg; 60-day release). A separate group of mice received a Placebo pellet as a control. Four weeks after pellet implantation, we collected blood from Placebo- and CORT-treated mice and used an enzyme-linked immunosorbent assay (ELISA) to quantify total plasma CORT levels (Fig.1A). As expected, there was a significant effect of treatment (Two-way ANOVA, *F*_(1,37)_=41.18, p<0.0001) on total plasma CORT levels (Fig.1B). We also observed a significant effect of sex on total plasma CORT levels (*F*_(1,37)_=11.78, p<0.01), and a significant effect of the interaction between treatment and sex (*F*_(1,37)_=17.25, p<0.001). Notably, we found that CORT pellet implant chronically elevated total plasma CORT in male mice only (Placebo Male vs CORT Male, Tukey’s multiple comparisons p<0.0001; CORT Male vs CORT Female, p<0.0001). The sex difference in total plasma CORT four weeks after CORT pellet implantation is consistent with previous studies in rats (27). However, a limitation of measuring total plasma CORT is that it includes both free and protein-bound CORT. Free CORT can cross the blood-brain barrier, while protein-bound CORT cannot (28,29). Corticosteroid binding globulin (CBG) is the primary protein in the blood that binds CORT and CBG is expressed at higher levels in females than males (30). Thus, we questioned if chronic CORT treatment decreased CBG in females, which would ultimately increase circulating levels of free CORT, even in the absence of changes in total levels. Indeed, chronic CORT treatment significantly decreased CBG levels in female mice (Fig.1C; Unpaired two-tailed t-test, p<0.05). Female mice continued to cycle through estrous normally, regardless of treatment (Fig.S1). We concluded that circulating levels of free CORT are likely elevated in both male and female mice after chronic CORT treatment with subcutaneous slow-release CORT pellets, albeit by different physiological mechanisms.

**Figure 1:**
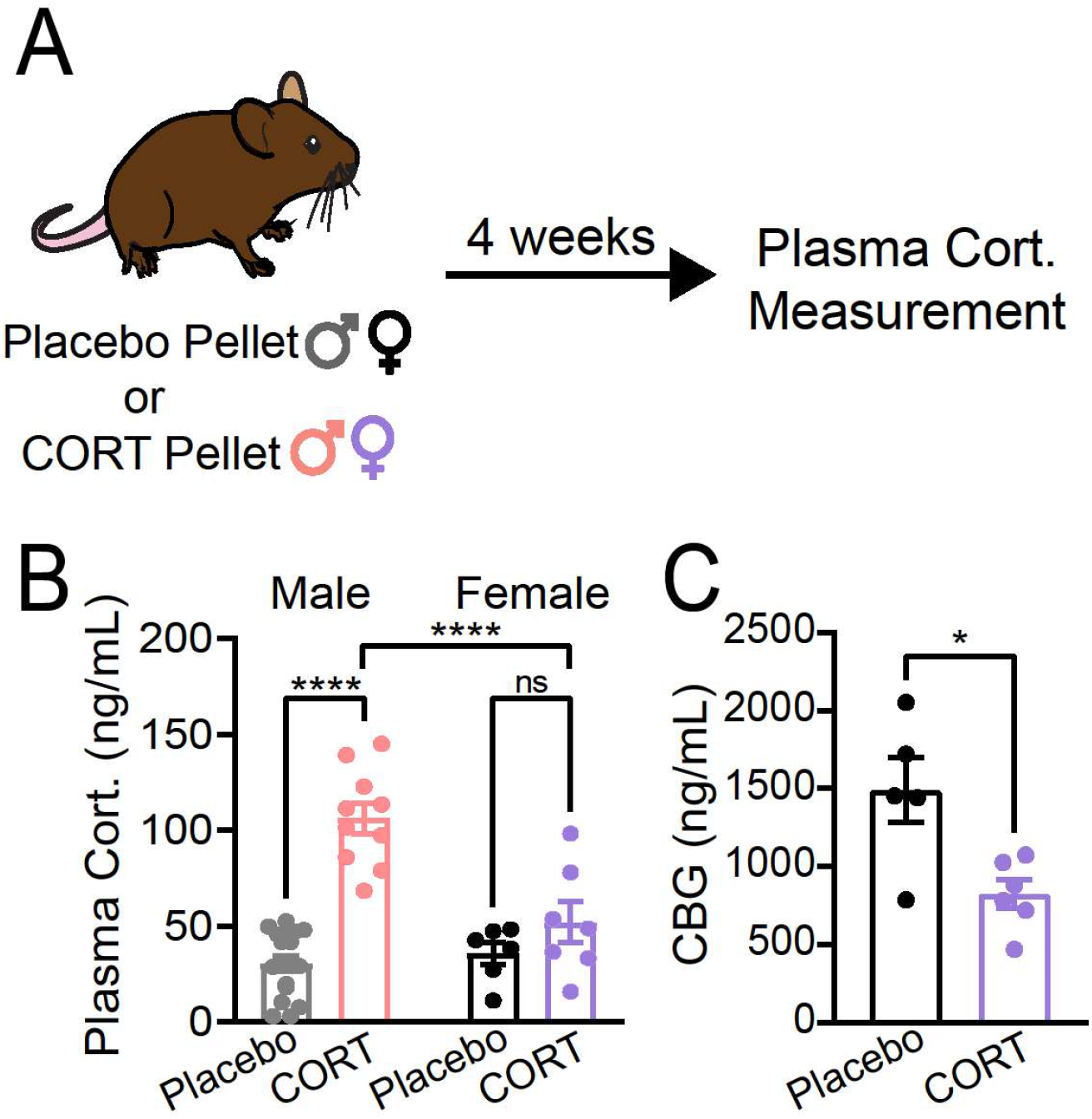
Chronic corticosterone treatment significantly increases plasma corticosterone in male mice and decreases corticosteroid binding globulin levels in female mice. A) Experimental timeline for pellet implantation and plasma corticosterone measurement. B) Plasma corticosterone (ng/mL) in male and female mice implanted with a placebo or corticosterone (35 mg; CORT) pellet. Two-Way ANOVA, main effect of treatment p<0.0001, main effect of sex p<0.01, main effect of treatment x sex interaction p<0.001, multiple comparisons ****p<0.0001. C) Plasma corticosteroid binding globulin (CBG; ng/mL) levels in placebo- and CORT-treated female mice (Unpaired two-tailed t-test *p<0.05). Each point represents an individual. Data presented as mean ± SEM.

### Chronic CORT treatment impairs reward-seeking in male and female mice

Chronic CORT treatment has previously been shown to impair reward-seeking behaviors in male mice (6,7). However, it was unclear what effect, if any, chronic CORT treatment would have on female mice. To assess reward-seeking behavior in both sexes, we used operant training. Four weeks after Placebo or CORT pellet implantation, mice began training on a fixed ratio-1 (FR-1) schedule, then advanced to FR-3 and FR-5 (Fig.2A). We found that CORT treatment significantly increased the number of days it took both sexes to acquire the FR-1 task (Fig.2B,F; Unpaired two-tailed t-test, p<0.05 male, p<0.0001 female), indicating a reward learning deficit caused by chronic CORT treatment. After FR-1 was acquired, impairments in reward-seeking behavior were observed in both sexes. Male mice displayed decreased rates of nosepoking (Fig.2C; significant effects of treatment [Two-way ANOVA *F*_(1,16)_=5.999, p<0.05], day of training [*F*_(8,128)_=41.28, p<0.0001], and an interaction between treatment and day of training [*F*_(8,128)_=2.226, p<0.05]). However, CORT treatment did not affect how often male mice interacted with the reward port across days of operant training (Fig.2D). Therefore, the decrease in the rates of rewards received by CORT-treated male mice during operant behavior was due to a reduction in reward seeking, but not reward retrieval (Fig.2E; Two-way ANOVA, *F*_(1,15)_=5.156, p<0.05).

**Figure 2:**
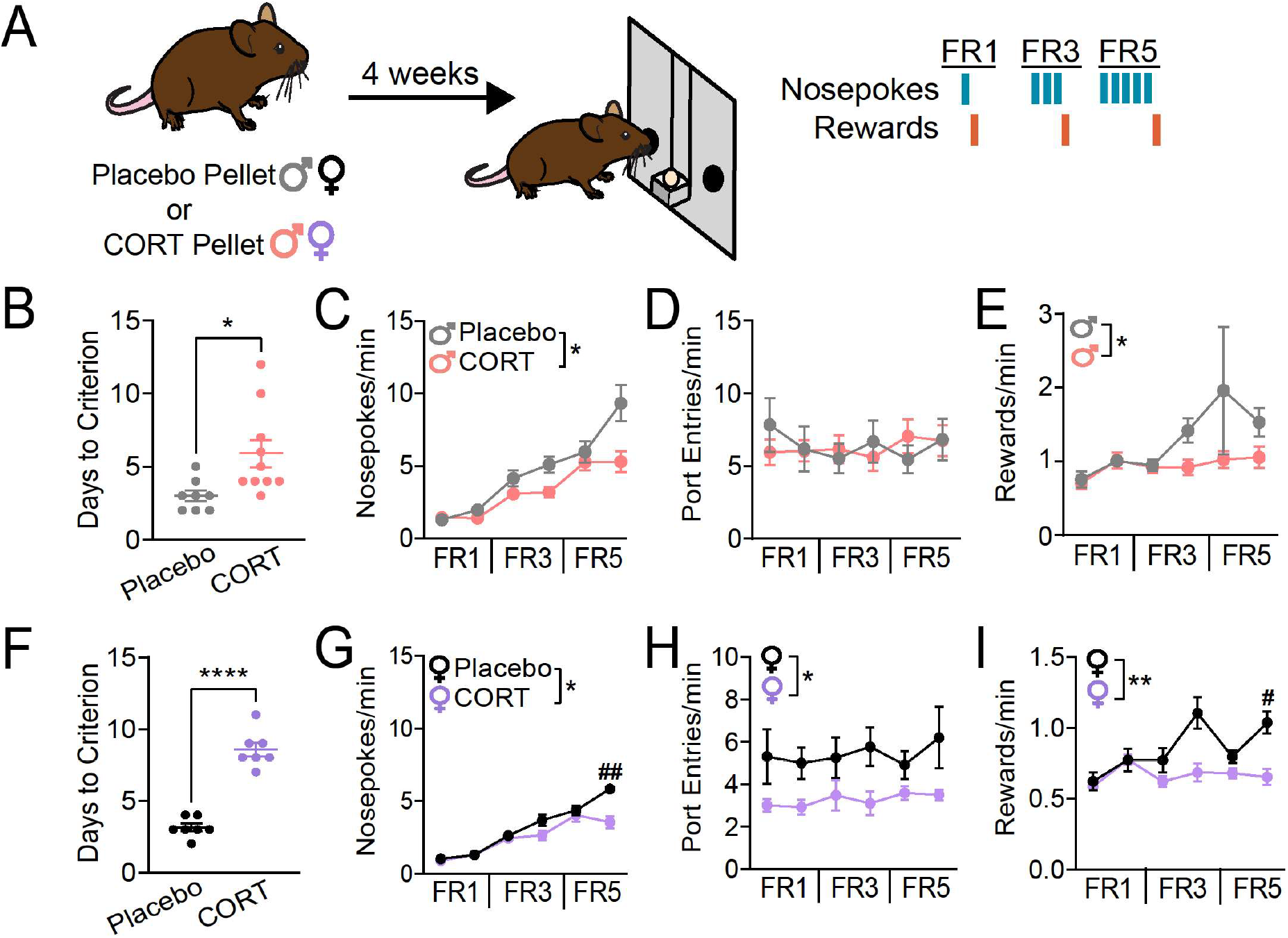
Chronic corticosterone treatment significantly decreases reward-seeking behavior in male and female mice and decreases consummatory behavior in female mice. A) Experimental timeline for pellet implantation and operant behavior paradigms. Right, schematic of fixed ratio (FR) paradigms. B) Days to acquire FR1 in male mice. Unpaired two-tailed t-test *p<0.05. Each point represents an individual. C) Nosepoking rates of Placebo-(N=8) and CORT-(N=10) treated male mice across operant behavior paradigms. Two-Way ANOVA, main effect of treatment *p<0.05, main effect of treatment x paradigm interaction p<0.001. D) Port entry rates of Placebo- and CORT-treated male mice across operant behavior paradigms. E) Reward rates of Placebo- and CORT-treated male mice across operant behavior paradigms. F) Two-Way ANOVA, main effect of treatment *p<0.05. G) Days to acquire FR1 in female mice. Unpaired two-tailed t-test ****p<0.0001. Each point represents an individual. H) Nosepoking rates of Placebo-(N=7) and CORT-(N=7) treated female mice across operant behavior paradigms. Two-Way ANOVA, main effect of treatment *p<0.05, main effect of treatment x paradigm interaction p<0.0001, multiple comparisons ##p<0.01. I) Port entry rates of Placebo- and CORT-treated female mice across operant behavior paradigms. Two-Way ANOVA, main effect of treatment *p<0.05. Reward rates of Placebo- and CORT-treated female mice across operant behavior paradigms. Two-Way ANOVA, main effect of treatment **p<0.01, main effect of treatment x paradigm p<0.01, multiple comparisons #p<0.05. Data presented as mean ± SEM.

Female mice displayed a significant effect of CORT treatment on nosepoking rates, with an interaction between training day and treatment (Fig.2G; significant effects of day of training [*F*_(3.267,39.20)_=87.47,p<0.0001], treatment [Two-way ANOVA *F*_(1,12)_=5.098, p<0.05], and an interaction between treatment and day of training [*F*_(5,60)_=7.294, p<0.0001]), driven by an effect on the last day of FR-5 (Sidak’s multiple comparisons p<0.01). In contrast to males, CORT treatment significantly decreased the port entry rate of female mice (Fig.2H; Two-way ANOVA, *F*_(1,12)_=5.062, p<0.05). Therefore, the decrease in the rate of rewards received by female mice stems primarily from a decrement in reward retrieval (Fig.2I; Two-way ANOVA, *F*_(1, 12)_=16.15, p<0.01), except on the last day of FR5 when it stems from the interaction between nosepoking and reward retrieval (Sidak’s multiple comparisons p<0.05).

### Chronic CORT treatment does not impair phasic dopamine transmission during reward-seeking

After observing an effect of CORT treatment on reward-seeking in both sexes, we questioned if CORT treatment was impairing phasic dopamine transmission in the striatum. To examine phasic dopamine transients in Placebo- and CORT-treated mice, we injected a virus encoding the fluorescent dopamine sensor, dLight1.3b (AAV9-CAG-dLight1.3b), into the NAcc and DMS of mice. We implanted a fiber optic over each injection site and recorded dLight1.3b transients during operant training. We found that CORT treatment had no effect on phasic dLight1.3b transients in the NAcc and DMS (Fig.S2), thus we turned our attention to measures of tonic dopamine activity in the striatum.

### Chronic CORT treatment decreases tissue dopamine content of the dorsomedial striatum (DMS) in female mice

To investigate whether chronic CORT treatment influenced the dopamine content of the striatum, we analyzed tissue samples from the NAcc and DMS of Placebo- and CORT-treated mice using high-performance liquid chromatography and electrochemical detection (HPLC-ECD) of dopamine. CORT treatment did not affect NAcc dopamine content in either sex (Fig.3B), but significantly decreased dopamine content of the DMS in female mice (Fig.3C; Unpaired two-tailed t-test, p<0.01). Therefore, although CORT has been reported to have acute effects on NAcc dopamine (31), we concluded that DMS dopamine is more sensitive to chronic CORT treatment, especially in females.

**Figure 3:**
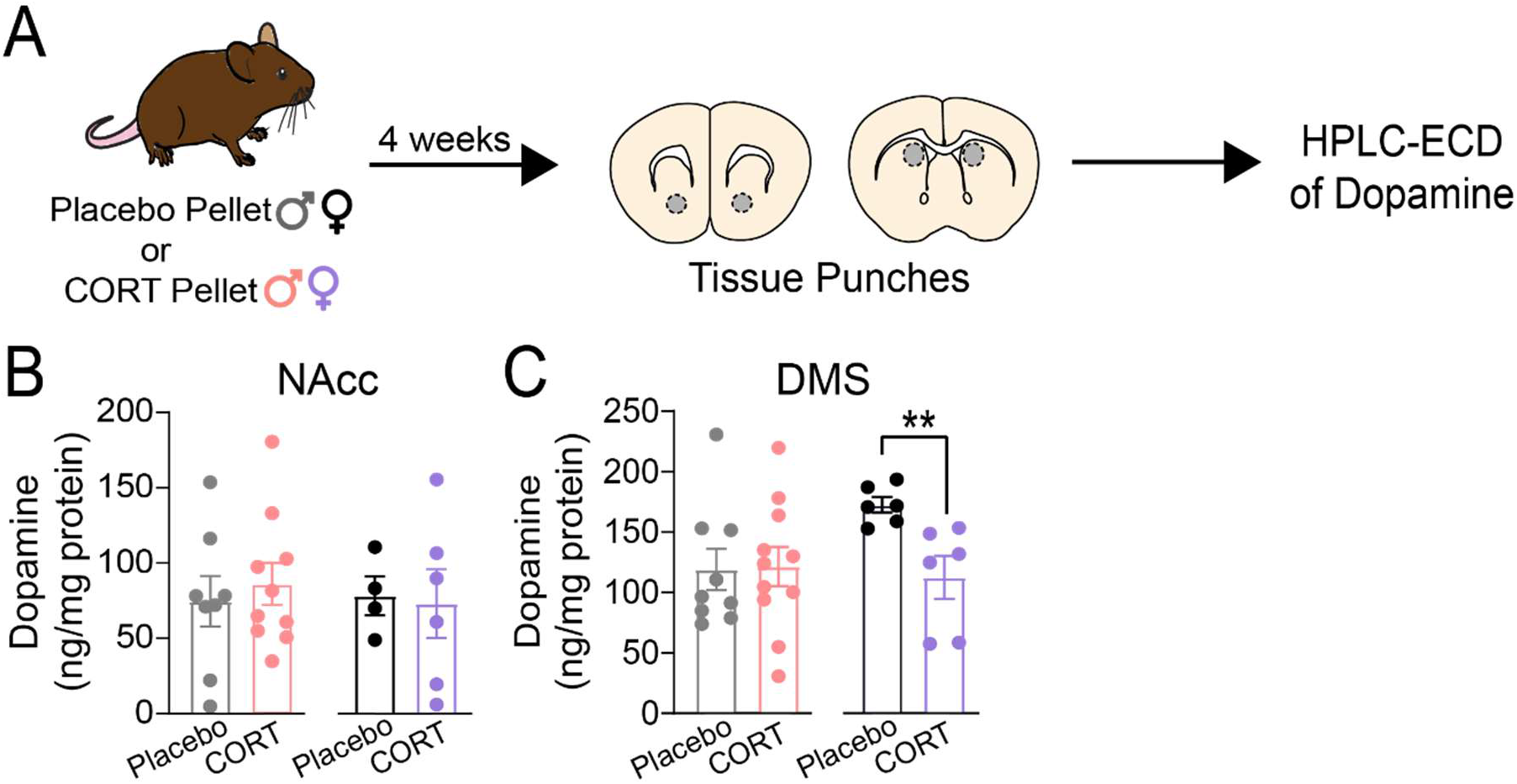
Chronic corticosterone treatment significantly decreases tissue dopamine content of the dorsomedial striatum (DMS) in female mice. A) Experimental timeline for pellet implantation, tissue punches, and HPLC-ECD of dopamine. B) Tissue dopamine content of the nucleus accumbens core (NAcc) of placebo- and CORT-treated male and female mice. C) Tissue dopamine content of the DMS of placebo- and CORT-treated male and female mice. Unpaired two-tailed t-test, **p<0.01. Each point represents an individual. Data presented as mean ± SEM.

### Chronic CORT treatment impairs dopamine transporter (DAT) function in the DMS of male mice

Another mechanism that can regulate levels of tonic dopamine in the striatum is modulation of dopamine transporter (DAT) function (16). By altering the decay rates of phasic dopamine transients, changes in DAT function can alter the timescale for integration of phasic dopamine signals, allowing or disallowing the buildup of tonic levels when dopamine neurons are active. When DAT function is impaired chronically, it can also lead to compensation in the dopamine system, altering the rate of synthesis of new dopamine (16). To investigate DAT function in mice treated chronically with CORT, we assayed dopamine dynamics in an *ex vivo* slice preparation. We injected a virus encoding the fluorescent dopamine sensor, dLight1.3b (AAV9-CAG-dLight1.3b), into the NAcc and DMS, and implanted Placebo or CORT pellets during the same surgery. Four weeks later, we prepared live striatal tissue sections and electrically evoked dopamine release using a bipolar stimulating electrode while simultaneously imaging dLight1.3b fluorescence (Fig.4A, Fig.S3). To mimic tonic and phasic dopamine neuron firing modes, we used a single stimulation pulse or a burst of 5 pulses at 20 Hz, respectively. We quantified the decay of dLight1.3b transients in response to electrical stimulation by calculating a ‘tau-off’ value and used it as a measure of the speed of extracellular dopamine clearance (23,32). CORT treatment did not significantly increase tau-off in male (Fig.4B,L) or female (Fig.4G,Q) mice in response to either one or five pulses at baseline in the DMS or in the NAcc (Fig.S4). However, the lack of change could be due to compensation for chronic DAT impairment. To elucidate how DAT was contributing to our measurements of dopamine clearance, we washed a DAT inhibitor, GBR12909 (1µM, ‘DATi’), onto the slice. We also tested the contribution of another monoamine transporter, Organic Cation Transporter 3 (OCT3) (33), to dopamine clearance by washing on an OCT3 inhibitor, normetanephrine (50µM; ‘OCTi’). OCT3 is a low-affinity, high-capacity non-specific monoamine transporter (34). Although OCT3 does not regulate synaptic dopamine levels as effectively as DAT, CORT has been demonstrated to bind OCT3 and inhibit monoamine reuptake, making it important to examine in our studies (33,35–37). When using either one or five stimulation pulses in the NAcc, we did not observe any significant effect of treatment on tau-off after DAT and OCT3 inhibition in either sex (Fig.S4). Using one stimulation pulse in the DMS, we did not observe significant differences in tau-off between Placebo- and CORT-treated mice after DAT and OCT3 inhibition (Fig.4D-F,I-K), although there was a trending effect of CORT treatment in male mice (Fig.4D, Two-way ANOVA, *F*_(1,16)_=3.636, p=0.07). In response to five pulses in the DMS, we found a significant effect of CORT treatment in male mice (Fig.4N-P; Two-way ANOVA, *F*_(1,14)_=8.566, p<0.05). DAT inhibition slowed dopamine clearance in Placebo-treated males as expected, but had no effect in CORT-treated males, indicating that DAT function is impaired in the DMS of CORT-treated males. Surprisingly, in females, we found no evidence for impairment of DAT or OCT3 function in the DMS following chronic CORT treatment (Fig.4S-U). CORT treatment did not affect the amplitude of dLight1.3b transients elicited in response to one or five stimulation pulses in the DMS of either sex (Fig.S5).

**Figure 4:**
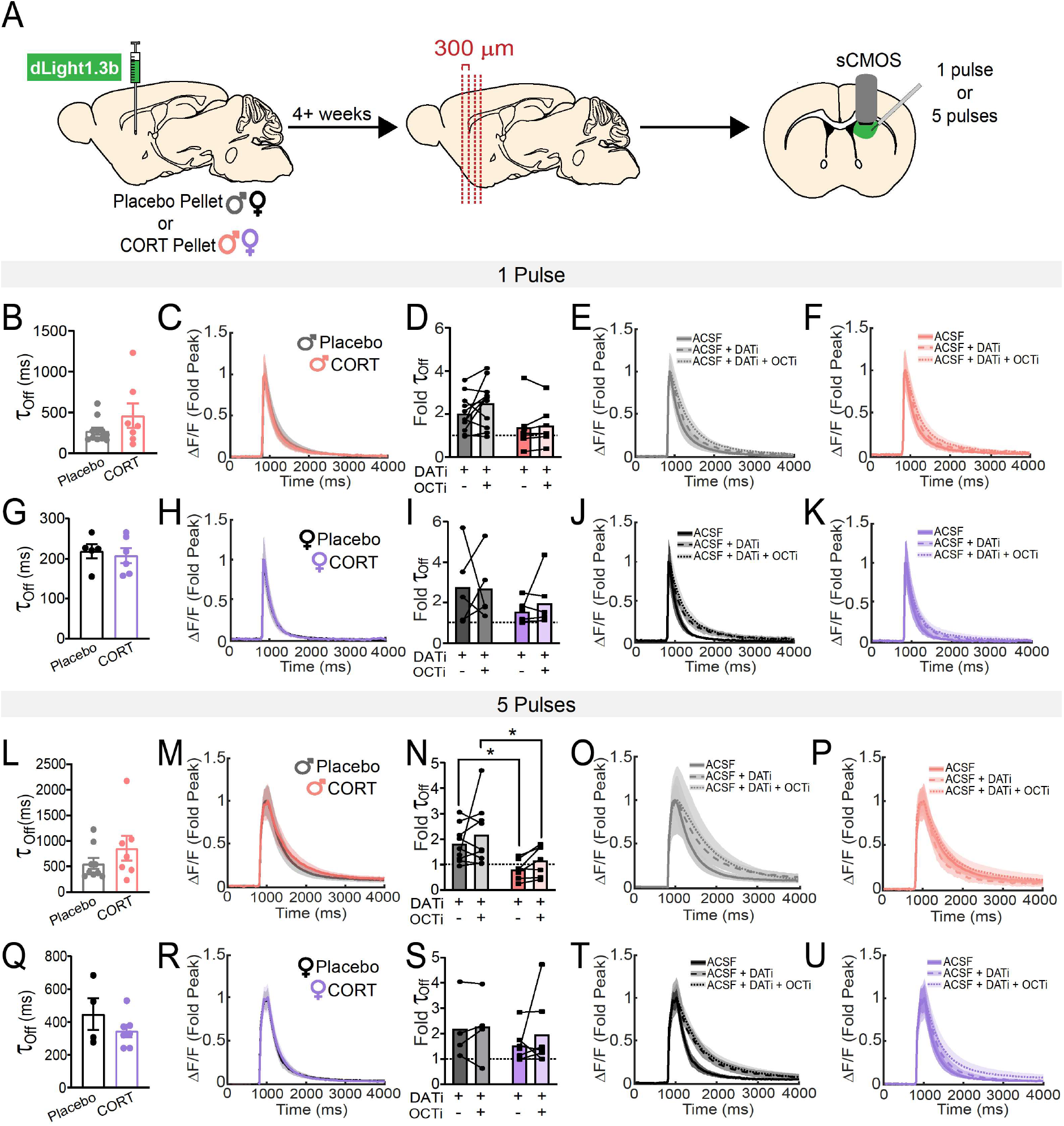
Chronic corticosterone treatment impairs *ex vivo* dopamine transporter (DAT) function in the dorsomedial striatum (DMS) of male mice. A) Experimental timeline for viral injection, pellet implantation, and slice imaging experiments. B) dLight1.3b fluorescence tau-off values after a single electrical stimulation of the DMS in acute tissue slices from male mice. C) Average dLight1.3b fluorescence traces, normalized to the peak of dLight1.3b fluorescence. after a single electrical stimulation of the DMS in acute tissue slices from male mice. D) Fold change of tau-off values of dLight1.3b fluorescence in the presence of inhibitors for the dopamine transporter (DATi) and organic cation transporter 3 (OCTi), normalized to tau-off values of dLight1.3b fluorescence in the absence of any transporter inhibitors, after a single electrical stimulation of the DMS in acute tissue slices from male mice. Two-Way ANOVA, trending effect of treatment p=0.07. E, F) Average dLight1.3b fluorescence traces, normalized to the peak of dLight1.3b fluorescence, after a single electrical stimulation of the DMS in acute slices from Placebo-(E) and CORT-(F) treated male mice, in the presence and absence of DATi and OCTi. G) dLight1.3b fluorescence tau-off values after a single electrical stimulation of the DMS in acute tissue slices from female mice. H) Average dLight1.3b fluorescence traces, normalized to the peak of dLight1.3b fluorescence, after a single electrical stimulation of the DMS in acute tissue slices from female mice. I) Fold change of tau-off values of dLight1.3b fluorescence in the presence of DATi and OCTi, normalized to tau-off values of dLight1.3b fluorescence in the absence of any transporter inhibitors, after a single electrical stimulation of the DMS in acute tissue slices from female mice. J, K) Average dLight1.3b fluorescence traces, normalized to the peak of dLight1.3b fluorescence, after a single electrical stimulation of the DMS in acute tissue slices from Placebo (J) and CORT-(K) treated female mice, in the presence and absence of DATi and OCTi. L) dLight1.3b fluorescence tau-off values after a 20 Hz, 5 pulse electrical stimulation of the DMS in acute tissue slices from male mice. M) Average dLight1.3b fluorescence traces, normalized to the peak of dLight1.3b fluorescence, after a 20 Hz, 5 pulse electrical stimulation of the DMS in acute tissue slices from male mice. N) Fold change of tau-off values of dLight1.3b fluorescence in the presence of DATi and OCTi, normalized to tau-off values of dLight1.3b fluorescence in the absence of any transporter inhibitors, after a 20 Hz, 5 pulse electrical stimulation of the DMS in acute tissue slices from male mice. Two-Way ANOVA, main effect of treatment p<0.05, multiple comparisons *p<0.05. O, P) Average dLight1.3b fluorescence traces, normalized to the peak of dLight1.3b fluorescence, after a 20 Hz, 5 pulse electrical stimulation of the DMS in acute tissue slices from Placebo-(O) and CORT-(P) treated male mice, in the presence and absence of DATi and OCTi. Q) dLight1.3b fluorescence tau-off values after a 20 Hz, 5 pulse electrical stimulation of the DMS in acute tissue slices from female mice. R) Average dLight1.3b fluorescence traces, normalized to the peak of dLight1.3b fluorescence, after a 20 Hz, 5 pulse electrical stimulation of the DMS in acute tissue slices from female mice. S) Fold change of tau-off values of dLight1.3b fluorescence in the presence of DATi and OCTi, normalized to tau-off values of dLight1.3b fluorescence in the absence of any transporter inhibitors, after a 20 Hz, 5 pulse electrical stimulation of the DMS in acute tissue slices from female mice. T, U) Average dLight1.3b fluorescence traces, normalized to the peak of dLight1.3b fluorescence, after a 20 Hz, 5 pulse electrical stimulation of the DMS in acute tissue slices from Placebo-(T) and CORT-(U) treated female mice, in the presence and absence of DATi and OCTi. Points represent the average of 2-3 sweeps from a single individual. Data presented as mean ± SEM.

To verify our *ex vivo* results, we designed an experiment to examine DAT function *in vivo* using fiber photometry. We injected a virus encoding dLight1.3b (AAV9-CAG-dLight1.3b) into the DMS of male and female mice and implanted a fiber optic in DMS for *in vivo* recording during behavior (Fig.S6). Four weeks later, mice were placed in an open field to collect baseline locomotor and dLight1.3b fluorescence data (Fig.5A). After ten minutes of baseline data collection, mice were injected with the DAT inhibitor, GBR12909 (20 mg/kg, i.p.), and returned to the open field for forty minutes. In mice with high DAT activity, injection of a DAT inhibitor should increase locomotion, with a concordant increase in extracellular dopamine in DMS (measured as a change in the dLight1.3b fluorescence area-under-the-curve (AUC)). In Placebo-treated male mice, we indeed observed robust increases in both locomotion and DMS dopamine in response to DAT inhibition. In CORT-treated male mice, we observed a blunting of locomotor activity in response to DAT inhibition accompanied by a blunted effect of DAT inhibition on DMS dopamine (Fig.5C-E). Comparing Placebo- and CORT-treated mice, we found significant effects of treatment (Two-way ANOVA, *F*_(1,17)_=6.776, p<0.05) and a trending effect of time (*F*_(2.344,39.86)_=2.835, p=0.06) on locomotor activity of male mice in response to DAT inhibition (Fig.5D). We also found significant effects of treatment (Two-way ANOVA, *F*_(1,9)_=5.418, p<0.05), time (*F*_(2.439,21.95)_=7.083, p<0.01), and the interaction between treatment and time (*F*_(49,441)_=2.979, p<0.0001) on dLight1.3b AUC after DAT inhibition in male mice (Fig.5E).

**Figure 5:**
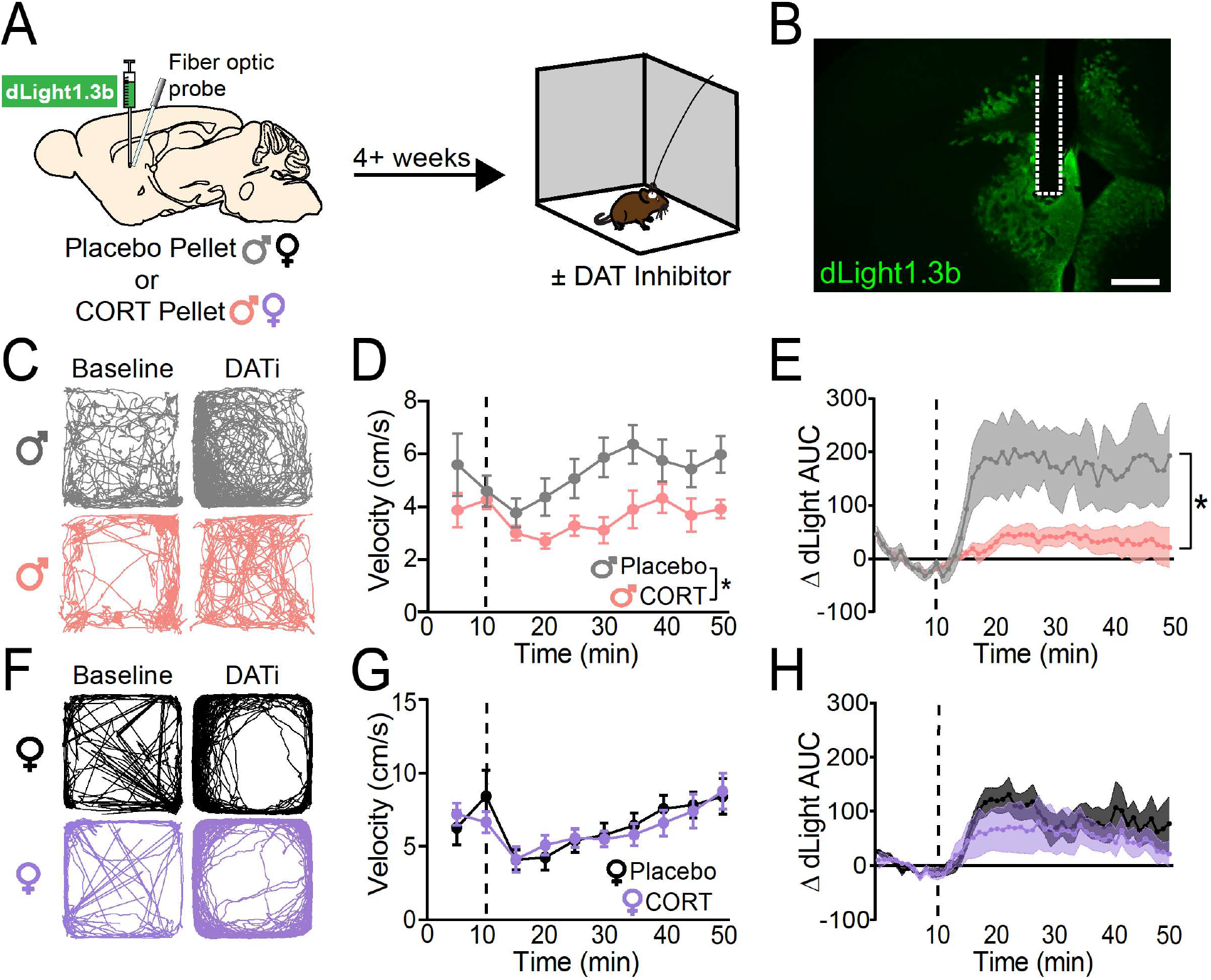
Chronic corticosterone treatment impairs *in vivo* dopamine transporter (DAT) function in the dorsomedial striatum (DMS) of male mice. A) Experimental timeline for viral injection, fiber optic implant, pellet implantation, and open field behavior. B) Representative image of dLight1.3b viral spread and fiber optic implantation site (outlined in dashed white line). Scale bar equals 500 micrometers. C) Representative activity traces of male mice (Placebo, top; CORT, bottom) during the ten-minute baseline period (‘Baseline’) and the last ten minutes of recorded activity after injection of the DAT inhibitor GBR12909 (20 mg/kg, i.p., ‘DATi’). D) Velocity of Placebo-(N=10) and CORT-(N=9) treated male mice, in averaged five-minute bins, before and after injection with the DAT inhibitor, GBR12909 (20 mg/kg, i.p.; injection time indicated by vertical dashed line). Two-Way ANOVA, main effect of treatment *p<0.05, trending effect of time p=0.06. E) Change in dLight1.3b area-under-the-curve (AUC) relative to the minute average of the ten-minute baseline period prior to injection with the DAT inhibitor, GBR12909 (20 mg/kg, i.p.; injection time indicated by vertical dashed line) in male mice. Two-Way ANOVA, main effect of treatment *p<0.05, main effect of time p<0.01, main effect of treatment x time interaction p<0.0001. Placebo N=5, CORT N=6. F) Representative activity traces of female mice (Placebo, top; CORT, bottom) during the ten-minute baseline period (‘Baseline’) and the last ten minutes of recorded activity after injection of the DAT inhibitor GBR12909 (20 mg/kg, i.p., ‘DATi’). G) Velocity of Placebo-(N=6) and CORT-(N=8) treated female mice, in averaged five-minute bins, before and after injection with the DAT inhibitor, GBR12909 (20 mg/kg, i.p.; injection time indicated by vertical dashed line). Two-Way ANOVA, main effect of time p<0.001. H) Change in dLight1.3b AUC relative to the average of the ten-minute baseline period prior to injection with the DAT inhibitor, GBR12909 (20 mg/kg, i.p.; injection time indicated by vertical dashed line) in female mice. Two-Way ANOVA, main effect of time p<0.01. Placebo N=5, CORT N=6. Data presented as mean ± SEM.

DAT inhibition could increase dLight1.3b AUC by increasing the decay constant of dLight1.3b transients, leading to a larger AUC per transient. Longer dopamine clearance times (indicated by higher decay constants) could then also slowly increase baseline dLight1.3b fluorescence, reflecting a slow buildup of tonic dopamine. Such an increase in baseline fluorescence would also contribute to an increase in dLight1.3b AUC following DAT inhibition. To look for these two effects and differentiate between them, we analyzed the decay constants of dLight1.3b transients recorded during open field behavior before and after DAT inhibition. We found that DAT inhibition increased both the decay time constants of *in vivo* dLight1.3b transients and led to a buildup of baseline dLight1.3b fluorescence (Fig.S7). We observed non-significant trends in which these effects were greater in Placebo-treated males than CORT-treated males (Fig.S7; Two-Way ANOVA, effect of treatment p=0.06 for decay constants, p=0.08 for baseline fluorescence). Thus, we concluded that the significant difference in DMS dLight1.3b AUC between Placebo- and CORT-treated males after DAT inhibition (Fig.5E) is the result of a *combined* effect on the decay of individual dLight1.3b transients and an increase in the baseline dLight1.3b fluorescence due to integration of slowly decaying transients.

In female mice, we did not observe differences between treatment conditions (Fig.5F-H). We observed a significant effect of time on locomotion (Fig.5G; Two-way ANOVA, *F*_(2.062,24.75)_=9.222, p<0.001) and on dLight1.3b AUC (Fig. 5H; Two-way ANOVA, *F*_(2.642,23.78)_=7.124, p<0.01) after DAT inhibition in both Placebo- and CORT-treated groups. We concluded that CORT treatment impairs DAT function in the DMS of male, but not female, mice. However, from these data it was not clear *how* CORT treatment was impairing DAT function.

### Chronic CORT treatment decreases phosphorylation of DAT at threonine-53

To assess how CORT treatment was impairing DAT function, we looked at DAT expression and post-translational modifications of DAT, which can affect reuptake activity (16,38,39). In particular, we were interested in examining phosphorylation at threonine-53, a known regulatory site (39–42). To determine if CORT treatment affected DAT expression or phosphorylation at threonine-53 in the DMS, we used western blot. We collected tissue punches from the DMS of Placebo- and CORT-treated male and female mice and fractionated the tissue homogenate to isolate membrane-bound proteins. We then probed for DAT and Thr53 phospho-DAT (pDAT). We found that CORT treatment had no effect on total membrane-bound expression levels of DAT in males or females (Fig.6C,D). However, CORT treatment significantly decreased pDAT in male mice (Unpaired, two-tailed t-test, p<0.05), but not female mice (Fig.6E,F). These results suggest that CORT treatment impairs DAT function in the DMS of male mice by decreasing phosphorylation of DAT at threonine-53, and further supports the conclusion that DMS DAT is unaffected by CORT treatment in female mice.

**Figure 6:**
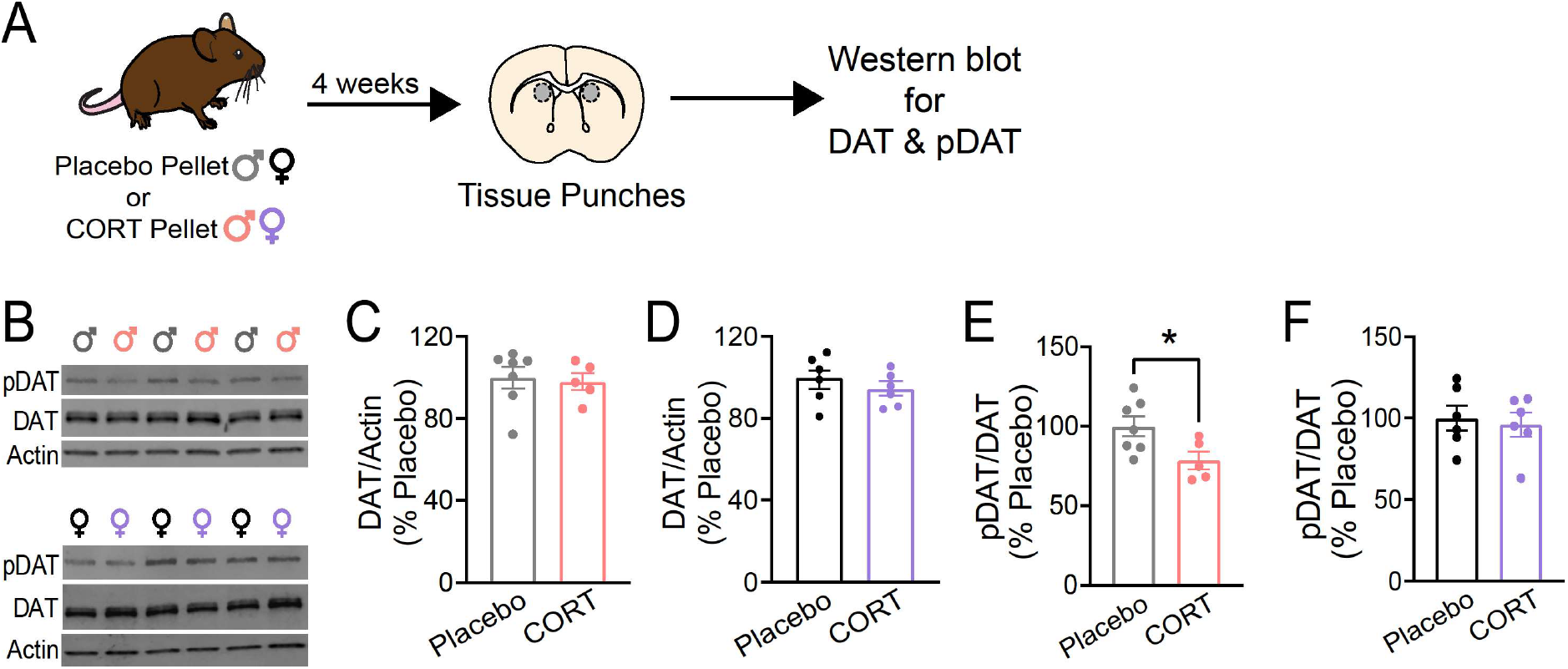
Chronic corticosterone treatment decreases phosphorylation of the dopamine transporter (DAT) at threonine-53 in the dorsomedial striatum (DMS) of male mice. .A) Experimental timeline for pellet implantation, tissue punches, and western blot experiments. B) Representative western blots for phosphorylated DAT at threonine-53 (‘pDAT’), DAT, and beta-actin (‘Actin’) from DMS tissue samples of male mice (top) and female mice (bottom). C) Membrane-bound DAT expression, normalized to Actin and plotted as a percent of Placebo expression, from DMS tissue samples of male mice. D) Membrane-bound DAT expression, normalized to Actin and plotted as a percent of Placebo expression, from DMS tissue samples of female mice. E) pDAT expression, normalized to DAT and plotted as a percent of Placebo expression, from DMS tissue samples of male mice. Unpaired two-tailed t-test *p<0.05. F) pDAT expression, normalized to DAT and plotted as a percent of Placebo expression, from DMS tissue samples of female mice. Each point represents a single individual. Data presented as mean ± SEM.

## Discussion

Previous studies in rodents had shown that chronic elevation of circulating CORT – a condition that can also occur in human MDD patients – impairs reward processing, but there was little mechanistic insight into *how* elevations in CORT impair reward processing. Further, preclinical literature had previously reported effects only in male rodents, yet humans with MDD are majority female. Here, we specifically set out to study both male and female mice and to identify mechanisms by which chronic CORT elevation might impact dopaminergic transmission, which is known to underlie reward-seeking behavior. We found that chronic CORT treatment impairs reward-seeking in both male and female mice (Fig.2C,G), but by sex-divergent mechanisms. In females, CORT treatment significantly decreases tissue dopamine content in the dorsomedial striatum (DMS; Fig.3C). In males, CORT treatment impairs dopamine transporter (DAT) function in DMS (Fig.4-6). Despite these differing mechanisms, we note that both males and females experienced changes in dopaminergic transmission specifically in DMS, tying dopaminergic function in this striatal subregion to the observed deficits in reward-seeking. Based on our results, we speculate that reward processing deficits observed in human MDD patients with dysregulated CORT may similarly be due to impaired DMS dopamine transmission, caused by distinct mechanisms in males and females. Our speculation is consistent with recent studies showing that individuals with MDD exhibit decreased DAT expression and tonic dopamine within the dorsal striatum (3,43). However, to address the aspect of our hypothesis dealing with sex differences, human data must be analyzed by sex. If the sex-divergent mechanisms by which DMS dopamine transmission is impaired in mice hold true for humans, this finding would suggest that medications for MDD should be tailored by sex.

Although numerous studies of male mice have reported deleterious effects of chronic CORT treatment on reward-seeking (6,7,44), to the best of our knowledge, our study is the first to report the effects of chronic CORT treatment in female mice. Notably, we observed a significant effect of chronic CORT treatment on reward consumption exclusively in females. Our results are surprising, given previous observations that CORT treatment has little effect on anxio-depressive behaviors in females (45,46). One reason we may have observed an effect of CORT treatment on reward-seeking in females is the route of administration of CORT. Previous studies primarily administered CORT in the drinking water (45,46), a route of administration that retains the circadian rhythm of plasma CORT levels. In contrast, pellet implantation abolishes circadian (and ultradian) CORT rhythmicity (47). Further studies are necessary to understand if abolishing CORT rhythmicity is necessary to observe effects of chronic CORT administration in females.

Critically, chronic CORT treatment did not affect dopaminergic transmission in the nucleus accumbens core (NAcc). Our findings are consistent with reports that adrenalectomy and CORT replacement do not affect extracellular dopamine levels in the NAcc (48). In contrast, our discovery that chronic CORT treatment primarily impacts dopaminergic transmission in the DMS is consistent with studies showing that DMS dopamine governs action-outcome learning, behavioral flexibility, and operant responding for rewards (11,12,49).

Our study adds to a growing body of literature demonstrating sex differences in DAT regulation in the dorsal striatum (39,50). Importantly, CORT’s lack of effect on DMS DAT function in females is likely not due to interaction with the estrous cycle, as it has been shown that the estrous cycle modulates dopamine reuptake in the ventral striatum, but not dorsal striatum (41,51). It remains unclear if restoration of DMS DAT function would ameliorate CORT-induced behavioral deficits in males. There is currently no intervention to precisely and exclusively manipulate DAT phosphorylation at threonine-53; however, generating the necessary genetic tools to achieve this manipulation will be a focus of future work.

Another critical open question is how chronic CORT treatment decreases DMS dopamine content in females, if not through a deficit in DAT function. We hypothesize that chronic CORT treatment could impair vesicular monoamine transporter 2 (VMAT2) function in females. VMAT2 exhibits greater activity in females than males, and an impairment in its function would decrease tonic dopamine levels in the DMS (52). We further hypothesize that female resistance to CORT-induced impairments in DAT function could be due to females’ higher capacity to sequester CORT in a protein-bound form (30) and/or their faster metabolism of CORT (53). Such differences could change how chronic CORT elevation impacts gene expression changes in dopamine neurons through pharmacodynamic differences in glucocorticoid receptor activation. Future studies are needed to address these possibilities in more detail and would be critical to identifying new sex-tailored drug targets for MDD patients with dysregulated CORT.

In sum, our studies suggest that a key mechanism underlying stress-induced deficits in reward processing is CORT’s impairment of DMS dopaminergic transmission. Our studies lay the groundwork for uncovering even more details about the relationships between CORT signaling and dopaminergic circuit function, such as regulation of DAT by endogenous circadian fluctuations in CORT. Given dopamine’s known neuromodulatory role in the dorsal striatum, it will also be interesting to investigate how chronic CORT treatment affects downstream DMS circuit function and corticostriatal plasticity to sustain the effects of chronic CORT treatment (54). The more we elucidate these pathways, identifying common mechanisms as well as sex differences, the more we will progress towards new therapeutic approaches for stress-related psychiatric disorders such as MDD.

## Acknowledgments

We thank the members of the Lerner laboratory for helpful discussions and critical feedback throughout the project. We thank Gates Palissery, Sean Pawelko, and Louis Van Camp for assistance with mouse breeding. We thank the Center for Comparative Medicine at Northwestern University for providing care for all mice used in these studies. We thank the Vanderbilt Neurochemistry Core for analysis of HPLC-ECD samples; the Neurochemistry Core is supported by the Vanderbilt Brain Institute and the Vanderbilt Kennedy Center. We thank the Alicia Guemez-Gamboa laboratory for advice and assistance with western blot experiments. We thank Drs. Joseph Bass, Yevgenia Kozorovitskiy, and Jones Parker for input and feedback on these studies. We additionally thank the Parker lab for assistance with dLight1.3b AUC analysis. This work was supported by a NARSAD Young Investigator Grant from the Brain and Behavior Research Foundation and an NIH New Innovator Award (DP2 MH122401) to T.N.L., and an NSF-GRFP Award (DGE-1842165) and NINDS DSPAN Award (F99NS130873-01) to A.L.H.

## Disclosures

The authors declare no competing interests.

## Notes

### Competing Interest Statement

The authors have declared no competing interest.

