## Supplementary Information & Figures for "Chronically elevated corticosterone impairs dopaminergic transmission in the dorsomedial striatum by sex-divergent mechanisms"

**Supplemental Methods**

<u>Methods</u>

*Tail Blood Sampling.*

Mice were gently restrained, and 30 µL of blood was sampled from the lateral tail vein. Plasma was separated out of blood by centrifugation at 4°C. Plasma samples were stored at -80°C prior to corticosterone and corticosteroid binding globulin (CBG) quantification.

*Corticosterone and Corticosteroid Binding Globulin (CBG) ELISAs (enzyme linked immunosorbent assays).*

Plasma corticosterone was quantified using a Cortiscosterone ELISA kit from Enzo Life Sciences, according to the manufacturer's instructions. Plasma CBG levels were quantified using a CBG ELISA kit from LifeSpan BioSciences, Inc., according to the manufacturer's instructions.

*Sample Preparation for High Performance Liquid Chromatography-Electrochemical Detection & Western Blot.*

After 4+ weeks after pellet implantation, mice were sacrificed by cervical dislocation and brains were flash-frozen in liquid nitrogen. Using a brain matrix, brains were sectioned into 1mm-thick sections and tissue punches of the nucleus accumbens core (NAcc) and dorsomedial striatum (DMS) were removed from tissue sections. Tissue punches were stored at -80°C. For HPLC-ECD, samples were sent to the Neurochemistry Core at Vanderbilt University for quantification of dopamine and its metabolites. For western blot, tissue punches were membrane fractionated using a Mem-PER Plus Protein Extraction kit from ThermoFisher, according to the manufacturer's instructions. A BCA Assay (ThermoFisher) was used to quantify protein content of samples. Samples were combined with Laemmli buffer and heated at 95°C for five minutes.

*High Performance Liquid Chromatography and Electrochemical Detection of Dopamine.*

Tissue Extraction: Tissues were kept frozen at -80°C and were held on dry ice prior to the addition of homogenization buffer in order to prevent degradation of biogenic amines. Tissues were homogenized, using a handheld sonic tissue dismembrator, in 100-750 µl of 0.1M TCA containing 0.01M sodium acetate, 0.1mM EDTA, and 10.5 % methanol (pH 3.8). Ten microliters of homogenate was used for the protein assay. The samples were then spun in a microcentrifuge at 10,000 g for 20 minutes. Supernatant was removed for HPLC-

ECD analysis. HPLC was performed using a Kinetix 2.6um C18 column (4.6 x 100 mm, Phenomenex, Torrance, CA USA). The same buffer used for tissue homogenization is used as the HPLC mobile phase.

Protein assay: Protein concentration in cell pellets was determined by BCA Protein Assay Kit (Thermo Scientific). Ten microliter tissue homogenate was distributed into 96-well plate and 200 µl of mixed BCA reagent (25 ml of Protein Reagent A is mixed with 500 µl of Protein Reagent B) was added. The plate was incubated at room temperature for two hours for the color development. A BSA standard curve was run at the same time. Absorbance was measured by a plate reader (POLARstar Omega), purchased from BMG LABTECH Company.

*Ex vivo dLight1.3b slice imaging analysis.*

Briefly, tau-off was calculated by isolating the decay of the dLight1.3b signal from a given sweep, fitting it with a single exponential, and solving for tau. To calculate the fold-effect size of the DAT inhibitor, GBR-12909, on dLight1.3b decay, tau-off values from the DAT inhibitor sweeps were calculated, then normalized to the average tau-off of dLight1.3b in the absence of the DAT inhibitor. The same was done for dLight1.3b traces acquired in the presence of the OCT3 inhibitor, Normetanephrine.

*In vivo dLight1.3b fiber photometry analysis.*

To calculate the baseline change in dLight1.3b fluorescence and the change in dLight1.3b decay after DAT inhibition *in vivo*, we normalized the dLight1.3b fluorescence by fitting it with the isosbestic fluorescence curve, then made peri-stimulus time histograms (PSTHs) of dLight1.3b transients in 10-minute intervals before and after injection with the DAT inhibitor, GBR12909. PSTHs included z-score data from 5 seconds prior to each dLight1.3b transient. We defined the 'baseline z-score' as the average z-score of first second of this 5 second pre-transient period. To calculate the decay constant of *in vivo* dLight1.3b transients, we normalized the average binned dLight1.3b decays to their peak z-score, then fit the decay of dLight1.3b transients with a double exponential. The timepoint in the normalized dLight1.3b decay where the z-score decreased to 36.8% of the fitted curve peak was defined as the decay constant.

*Statistical Analysis.*

All statistical analyses were completed using GraphPad Prism. P-values less than 0.05 were considered statistically significant.

**Supplemental Figures**

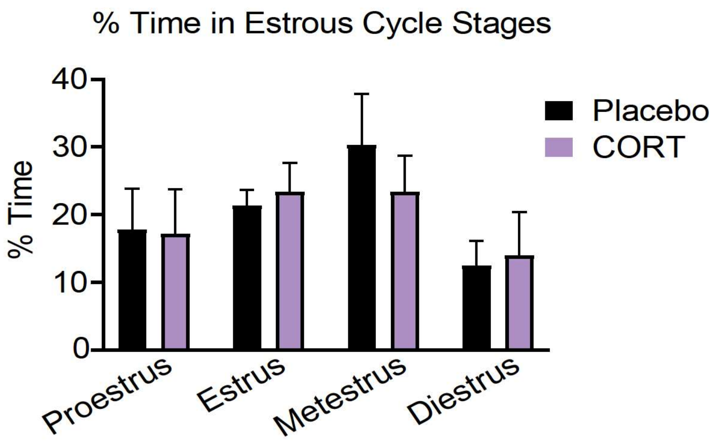


**Supplemental Figure 1: Chronic corticosterone treatment does not affect estrous cyclicity of female mice.**

Percent of time spent in each phase of the estrous cycle over the course of eight days of sampling. Placebo N=7, CORT N=8.

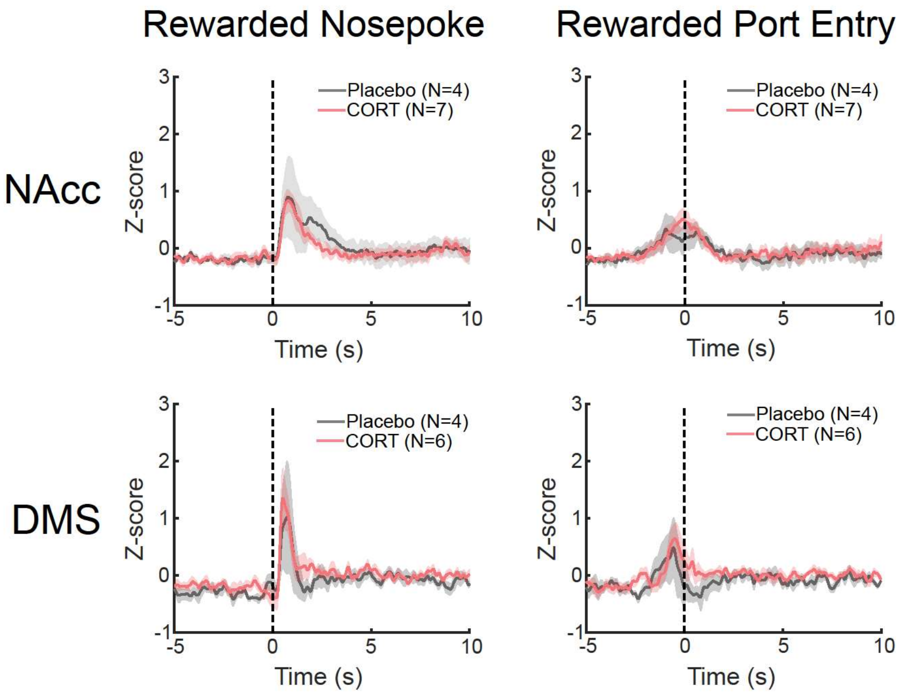


**Supplemental Figure 2: Chronic corticosterone treatment does not affect phasic dopamine transmission during operant training.**

Peri-stimulus time histograms of dLight1.3b fluorescence recorded from nucleus accumbens core (NAcc; top) and dorsomedial striatum (DMS; bottom) aligned to the time of a rewarded nosepoke (left) and rewarded port entry (right) during FR-5 training. Data from male mice only. Data presented as mean ± SEM.

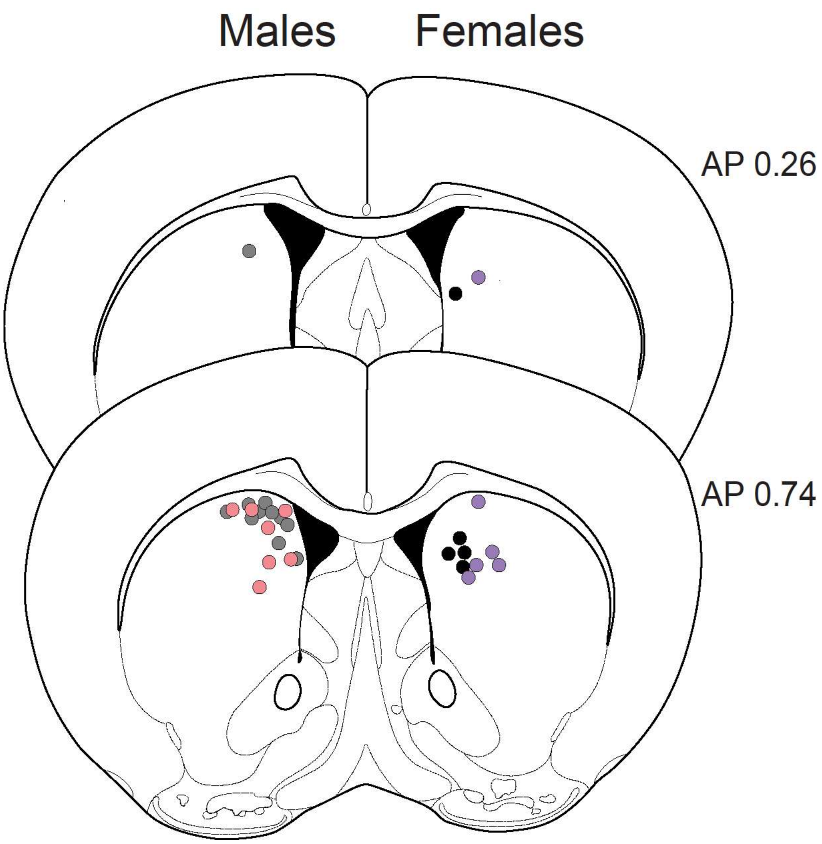


**Supplemental Figure 3: *Ex vivo* dLight1.3b recording sites in the dorsomedial striatum (DMS).** Recording sites from Placebo-treated males (gray), CORT-treated males (pink), Placebo-treated females (black), and CORT-treated females (purple). AP coordinate indicates slice position relative to bregma.

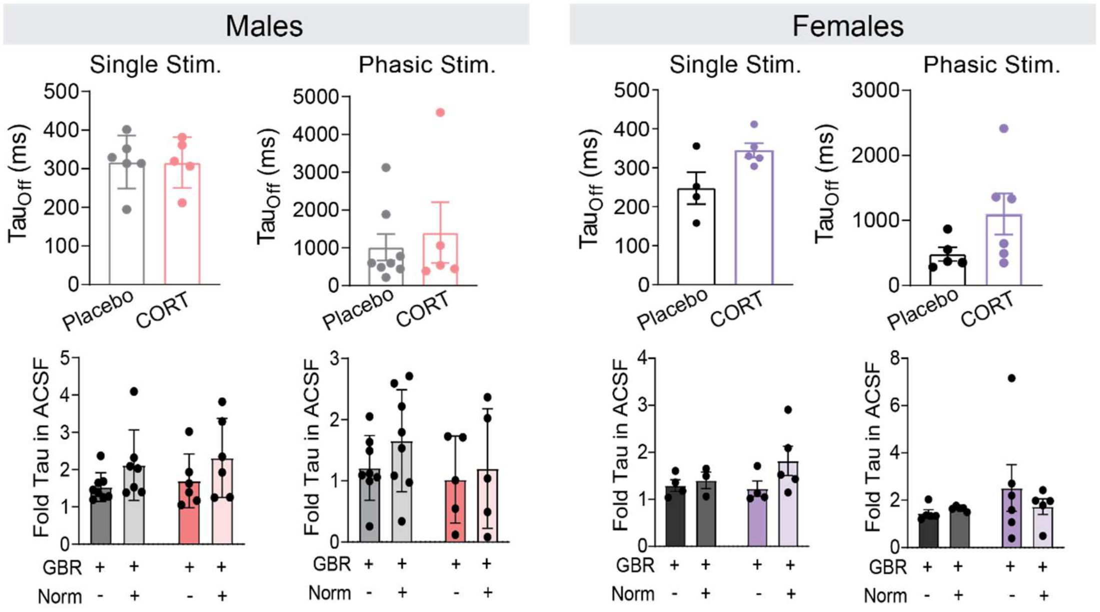


**Supplemental Figure 4: Chronic corticosterone treatment does not affect *ex vivo* dopamine transporter (DAT) function in the nucleus accumbens core (NAcc) of either male or female mice.** Top: tau-off of dLight1.3b signal in the NAcc after a single or phasic stimulation, in males (left) and females (right). Bottom: fold change of tau-off values of dLight1.3b fluorescence in the presence of DATi and OCTi, normalized to tau-off values of dLight1.3b fluorescence in the absence of any transporter inhibitors, after a single or phasic stimulation in the NAcc of males (left) and females (right).

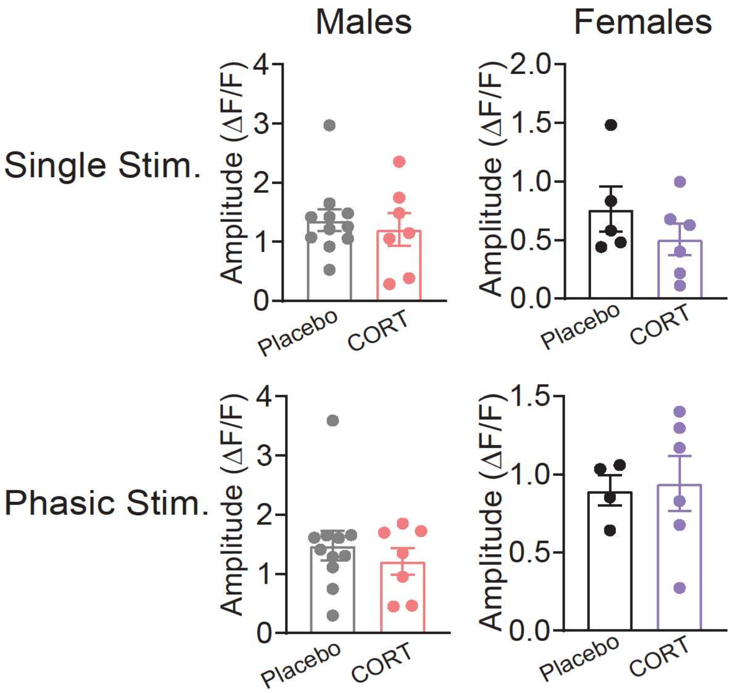


**Supplemental Figure 5: Chronic corticosterone treatment does not affect amplitude of electrically-evoked dLight1.3b fluorescence in the dorsomedial striatum (DMS) of male or female mice.**

Amplitudes of electrically-evoked dLight1.3b events, normalized to baseline fluorescence, in the DMS of male (left) and female (right) mice, in response to a single electrical stimulation (top) or phasic electrical stimulation (bottom).

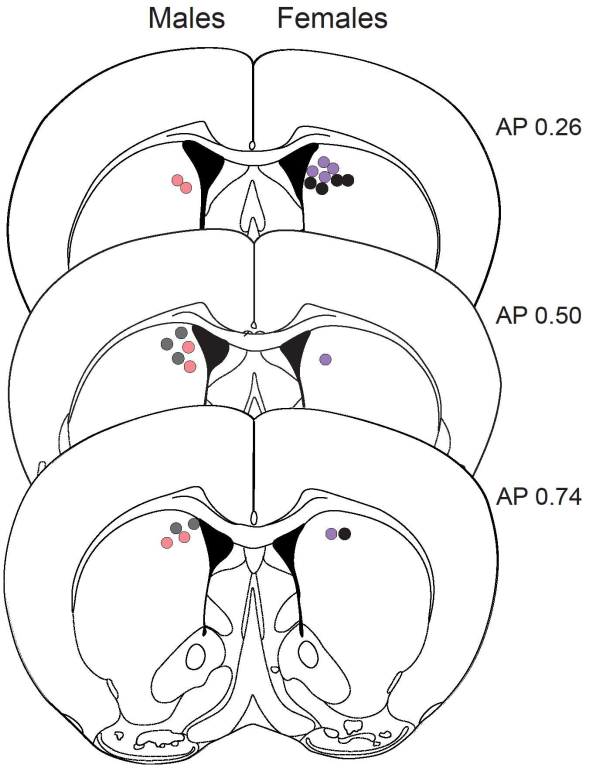


**Supplemental Figure 6: Fiber optic probe implant sites.** Terminal sites of fiber optic probes in Placebo-treated males (gray), CORT-treated males (pink), Placebo-treated females (black), and CORT-treated females (purple). Each point represents one individual. AP coordinate indicates slice position relative to bregma.

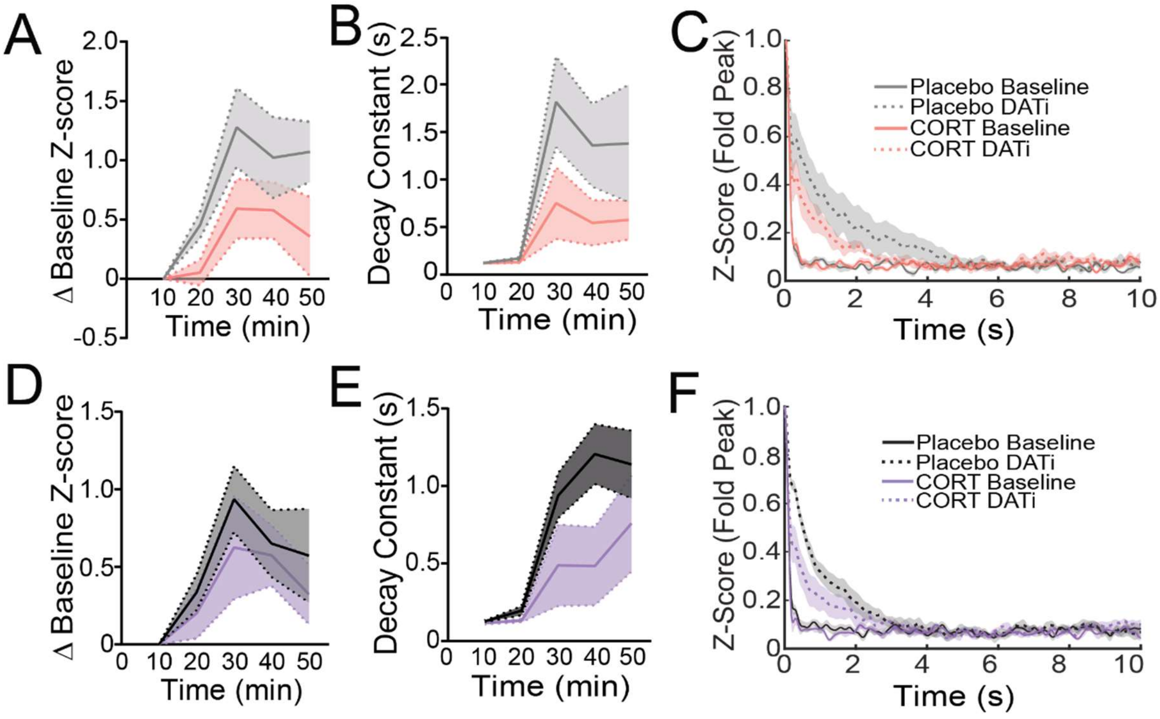


**Supplemental Figure 7: Chronic corticosterone treatment tends to blunt increased baseline fluorescence and decay constant of dLight1.3b after DAT inhibition in the dorsomedial striatum (DMS) of male mice.**

A) Change in baseline z-score of dLight1.3b fluorescence in the DMS after DAT inhibition in males (Two-Way ANOVA, main effect of time $p<0.01$, trending effect of treatment $p=0.08$). Placebo N=5, CORT N=6.

B) Decay constant of dLight1.3b in the DMS before and after DAT inhibition in males (Two-Way ANOVA, main effect of time $p<0.001$, trending effect of treatment $p=0.06$).

C) Average dLight1.3b decay during the 10 minutes before DAT inhibition ('Baseline') and during the last 10 minutes of recorded dLight1.3b fluorescence after DAT inhibition ('DATi') in the DMS of male mice.

D) Change in baseline z-score of dLight1.3b fluorescence in the DMS after DAT inhibition in females (Two-Way ANOVA, main effect of time $p<0.01$). Placebo N=5, CORT N=6.

E) Decay constant of dLight1.3b in the DMS before and after DAT inhibition in females (Two-Way ANOVA, main effect of time $p<0.001$).

F) Average dLight1.3b decay during the 10 minutes before DAT inhibition ('Baseline') and during the last 10 minutes of recorded dLight1.3b fluorescence after DAT inhibition ('DATi') in the DMS of female mice.

Data presented as mean ± SEM.
